# Fractional response analysis reveals logarithmic cytokine responses in cellular populations

**DOI:** 10.1101/2020.12.08.413468

**Authors:** Karol Nienałtowski, Rachel E. Rigby, Jarosław Walczak, Karolina E. Zakrzewska, Jan Rehwinkel, Michał Komorowski

## Abstract

Although we can now measure single-cell signaling responses with multivariate, high-throughput techniques our ability to interpret such measurements is still limited. Even interpretation of dose-response based on single-cell data is not straightforward: signaling responses can differ significantly between cells, encompass multiple signaling effectors, and have dynamic character. Here, we use probabilistic modeling and information-theory to introduce fractional response analysis (FRA), which quantifies changes in fractions of cells with given response levels. FRA can be universally performed for heterogeneous, multivariate, and dynamic measurements and, as we demonstrate, uncovers otherwise hidden patterns in single-cell data. In particular, we show that fractional responses to type I interferon in human peripheral blood mononuclear cells are very similar across different cell types, despite significant differences in mean or median responses and degrees of cell-to-cell heterogeneity. Further, we demonstrate that fractional responses to cytokines scale linearly with the log of the cytokine dose, which uncovers that cellular populations are sensitive to fold-changes in the dose, as opposed to additive changes.

## Main text

Many studies of signaling systems involve examining how the intensity of a stimulus, e.g., cytokine dose, translates into the activity of signaling effectors, e.g., transcription factors^1–7^. This is usually done by exposing cells to a range of doses and measuring responses either in bulk or at the single-cell level. Results of such experiments are then represented and interpreted in terms of dose-response curves. The standard dose-response curve depicts how the mean, median, or a characteristic of choice, changes with the increasing dose, and provides a basic, first-order model of how a signaling system operates. Several aspects of cellular signaling are difficult to analyze using conventional approaches. For example, signaling responses can differ significantly between cells, encompass multiple signaling effectors, and are dynamic. First, outwardly very similar cells exposed to the same stimulus exhibit substantial cell-to-cell heterogeneity^8–12^. Therefore the same mean/median response can result from a small fraction of strongly responding cells or a significant fraction of weakly responding cells^1,2,13^. Second, the highly interconnected architecture typical for mammalian signaling usually results in a single stimulus activating several primary signaling effectors or downstream genes^14–19^. For example, effectors of type I interferons (IFNs) include six members of the signal transducer and activator of transcription family (STAT)^20^, which are activated with different sensitivities at different doses. Therefore, the description of dose-response in terms of an individual signaling effector is incomplete^21^. Third, live-cell imaging experiments demonstrated that the dose may not only alter the response at a single time-point but can control temporal profiles of signaling responses^22,23^. For instance, low doses of TNF-*α* may induce one peak of nuclear factor-*κ*B (NF-*κ*B) signaling activity, whereas higher doses may induce additional peaks^7,24^. Besides, the dose may control the onset, shut off, amplitude, or, in principle, any other characteristics of the responses^25–28^. Overall, conventional dose-response curves do not capture the inherent complexity of single-cell high-throughput data, and an alternative approach is required. We have used probabilistic modeling and information-theory to develop a different analytic framework, fractional response analysis (FRA), involving fractional cell counting, which is capable of deconvoluting the behavior of single cells.

## Results

### Conventional dose-response analysis does not capture complex data

To demonstrate the need and utility of FRA we studied type I interferon signaling in human peripheral blood mononuclear cells (PBMCs), a system involving multiple signaling effectors, cell-to-cell heterogeneity, and several cell types. Dose-responses to the type I interferon variant IFN-*α*2a were analyzed via whole-cell tyrosine phosphorylation levels of effector proteins STAT1, STAT3, STAT4, STAT5, and STAT6 (pSTATs), using mass cytometry (CyTOF). Cells were collected from a healthy donor, and measurements were performed 15 minutes after IFN-*α*2a stimulation, the time of maximal response (Fig. S1). Along with signaling effectors, 26 phenotypic markers were measured to allow for identification of several cell types, including B-cells, CD4+ T-cells, CD8+ T-cells, natural killer (NK) cells, and CD14+ monocytes^29–31^. Such data are typically analyzed using t-SNE plots to visualize multiple cell types and signaling effectors^30,31^ (Fig. 1A-B, Fig. S2). However, a single t-SNE plot represents responses in terms of one signaling effector only, and plots do not provide any quantitative information regarding the function of the signaling systems.

**Figure 1.**
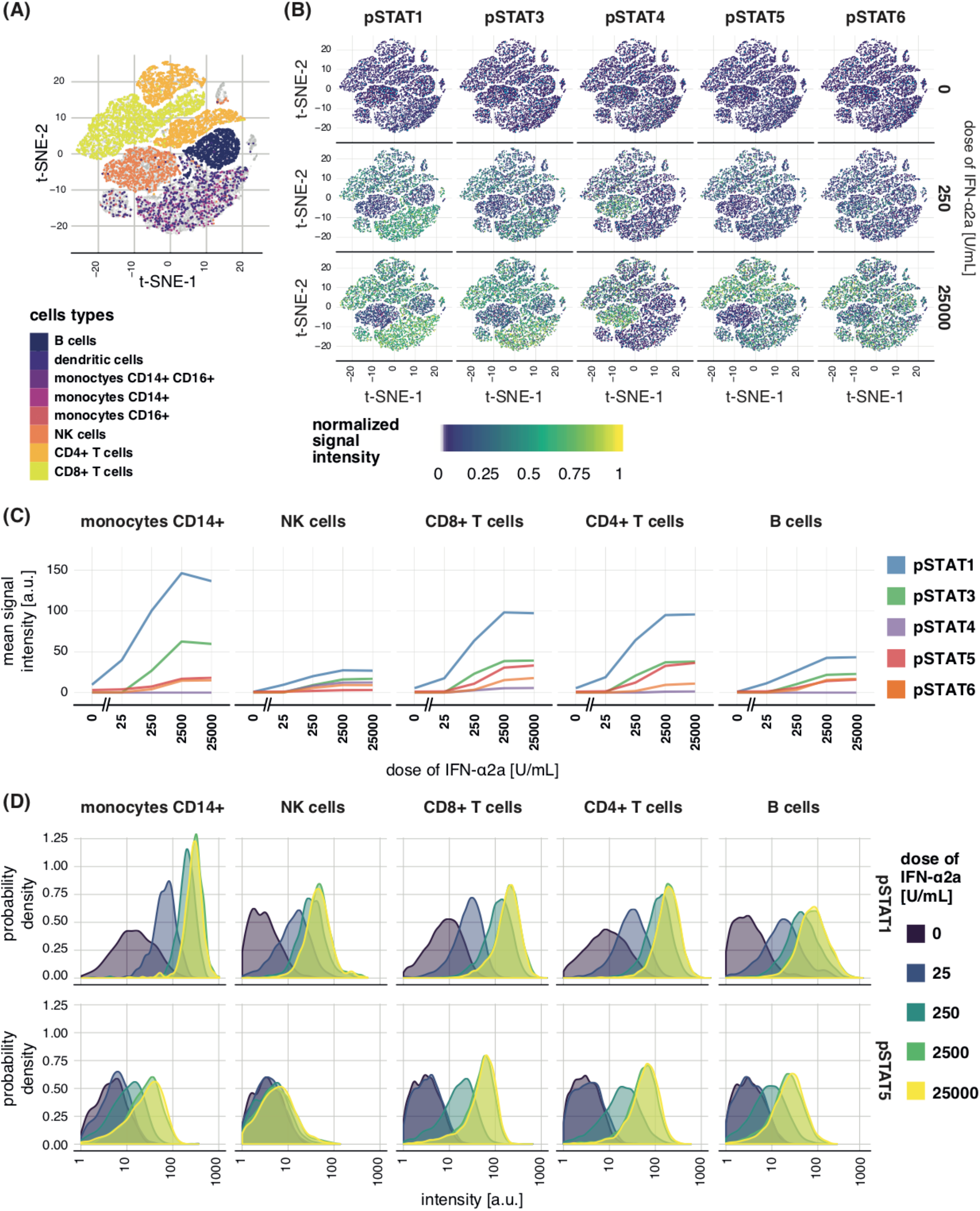
Dose-responses to IFN-*α*2a in PBMCs. **(A)** t-SNE plots constructed based on phenotypic markers. Cell types are encoded by color and each dot represents a single cell. **(B)** t-SNE plots of whole-cell pSTATs levels 15 min after stimulation with two selected doses of IFN-*α*2a as well as in unstimulated cells. Positions of dots corresponding to single-cells are the same as in panel (A) allowing cell type identification. Color of each dot represents normalized (0 for minimum and 1 for maximum) mass cytometry signal. Analogous t-SNE plots for all considered doses are show in Fig. S2. **(C)** Mean pSTATs levels in five cell types as a function of dose calculated from mass cytometry signals of single cells. **(D)** Distributions of responses in five cell types after stimulation with different doses of IFN-*α*2a in terms of pSTAT1 (top row) and pSTAT5 (bottom row) as measured with mass cytometry. The shown probability density is proportional to the frequency of cells with given level of the pSTAT. Value of the probability density is proportional to the frequency of cells with given response levels. Distributions of other pSTATs are show in Fig. S3. Different doses correspond to different colors.

Typically, to obtain quantitative characteristics of the dose-response, mean responses and response distributions of individual signaling effectors are plotted. Following this conventional strategy, mean levels of pSTATs in B-cells, CD4+ T-cells, CD8+ T-cells, NK cells, and CD14+ monocytes, are calculated (Fig. 1C, Fig. S3). Mean responses revealed that different STATs reached different maximal phosphorylation levels in different cell types, and response distributions indicated that the variability of responses can vary between cell types. Overall, however, analysis using conventional techniques (Fig. 1A-D), did not reveal any apparent pattern in functioning of the signaling system. The failure to observe regularities resulted largely from the complexity of the system and of the data.

### Fractional response curves

To deconvolute single-cell dose-response data, we first introduced the fractional response curve (FRC) that quantifies fractions of cells that exhibit different responses to a change in dose, or any other experimental condition. To illustrate the concept, we considered a simple hypothetical example involving one signaling effector and three doses, although the approach extends to a general multivariate scenario. Response distributions to three doses, *x*_1_, *x*_2_, *x*_3_, which can be interpreted as control, intermediate and high dose, are shown in Fig. 2A. When dose 1 was considered alone, fractions of cells with all possible responses sum up to 1 (Fig. 2B). Therefore, we defined the value of the FRC for dose 1 to be 1, and write *r*(*x*_1_)= 1. We then asked what fraction of the cellular population exhibits different responses after the change from dose 1 to dose 2. The fraction of cells exhibiting different responses is equivalent to the overall increase in the frequency of responses (Fig. 2C, green region). The overall fractional increase, denoted as Δ*r*, is calculated as the area of the green region, and Δ*r* = 0.31, represents the 31% of the cellular population exhibiting different responses due to dose increase. Therefore, we defined the value of the FRC for dose 2 to be the sum of the previous value and the fractional increment, *r*(*x*_2_)= *r*(*x*_1_) + Δ*r =* 1.31. When dose 3 was considered, the fraction of cells that exhibited different responses is again equivalent to the overall increase in the frequency of different responses, now compared to the two lower doses (Fig. 2D). As before, the overall increase, Δ*r*, is equivalent to the area of the yellow region (Fig. 2D), with Δ*r* = 0.74, representing 74% of cells stimulated with dose 3 exhibiting responses different to populations stimulated with lower doses. Again, the value of the FRC for dose 3 was defined as the sum of previous value and the fractional increment, *r*(*x*_3_)= *r*(*x*_2_) + Δ*r =* 2.05. Changes in the FRC show what fraction of cells exhibit different responses due to the dose increase (see *Methods* for a formal definition). Adding subsequent fractional increments, Δ*r*, leads to the value of FRC expressed in terms of the cumulative fraction of cells that exhibit different responses due to dose change. Besides, the sum of the dose-to-dose increments records the number of distinct response distributions that were experimentally observed, which provides the second interpretation of the FRC. Precisely, for dose 1 considered alone, a single response distribution was observed, *r*(*x*_1_)= 1. Dose 2 added 31% of a distinct distribution, and *r*(*x*_2_)= 1.31 (the gray area, Fig. 2D). Similarly, accounting for all three doses we had 2.05 distinct response distributions (the gray area, Fig. 2E). The number of distinct response distributions induced by changing dose quantifies the number of programmed responses of a cellular population, which appears to provide relevant, yet, so far, unexplored, quantitative characteristics of signaling systems (Fig. S4).

**Figure 2.**
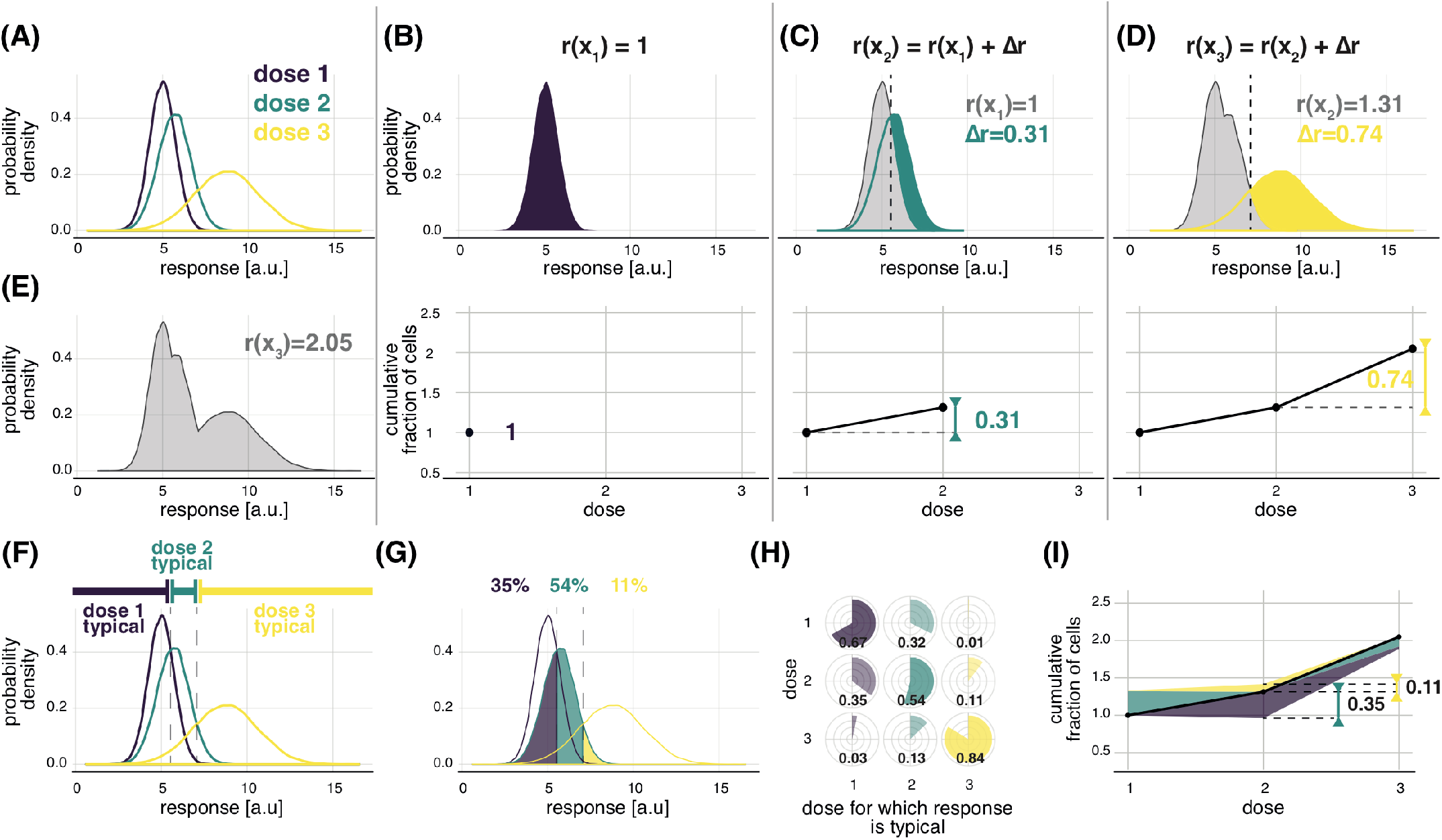
Fractional response analysis. **(A)** Hypothetical response distributions to three different doses encoded by colors. Distributions are represented as probability density, which is proportional to the frequency of cells with a given response level. **(B-D)** Quantification of the fraction of cells that exhibit different responses due to dose increase, Δ*r*, and constriction of FRC, for responses presented in (A). Each panel from B to D correspond to subsequent changes in dose. The color regions mark the overall increase in frequency due to considering the dose marked by the color. The area of the colored region quantifies Δ*r*. The value of the FRC for each dose is obtained by adding the increment, Δ*r*. **(E)** Quantification of the number of distinct distributions induced by the three considered doses. **(F)** Dose-typical responses for the response distributions of (A). **(G)** Dissection of the responses to dose 2 into responses typical to any of the three doses. The fraction of cells typical to a given dose is marked with the corresponding color. The surface area of each color quantifies the typical fraction. **(H)** The fractions of cells stimulated with one dose (rows) with responses typical to any of the doses (columns). **(I)** The FRC together with the bands representing cell-to-cell heterogeneity as quantified in (K). For each reference dose (x-axis), the fractions of cells stimulated with the reference dose that exhibit responses typical to other doses can be plotted in the form of color bands around the curve. The color encodes the dose a given fraction refers to. The height of the band marks the size of the fraction (y-axis). Fractions corresponding to doses higher than the reference dose are plotted above the curve, whereas to doses lower than the reference dose below the curve.

The FRC has a rigorous mathematical interpretation in terms of Rényi information, which, broadly speaking, counts probability distributions corresponding to outputs of a communication system (see *Supplementary Material, SM)*. The FRC can be calculated for any type of signaling data, i.e., arbitrary number of signaling effectors, time points of measurements, doses, or other experimentally varied parameter (see *Methods*).

### Fractional cell-to-cell heterogeneity

The FRC quantifies fractions of cells that exhibit different responses due to dose change but does not quantify cell-to-cell heterogeneity: it does not show what fraction of cells exposed to one dose exhibits responses in the range characteristic for other doses. Therefore, within FRA, we propose to augment the FRC with quantification of the overlaps between distributions corresponding to different doses. We call a given response as typical for a given dose if it is most likely, i.e., most frequent, to arise for this specific dose compared to all other doses. In the hypothetical example, low responses are most likely, and therefore typical, for dose 1, intermediate responses are typical for dose 2, and high responses for dose 3 (Fig. 2F). We can then divide responses to a given dose into responses typical for any dose. For instance, for dose 2, 35% of cells have responses typical for dose 1, 54% typical for dose 2, and 11% typical for dose 3 (Fig. 2G). The results, presented as pie-charts, can be shown in a matrix as the fraction of cells stimulated with one dose (rows) that has responses typical for other doses (columns) (Fig. 2H). This pie-chart partitioning can be plotted along the FRC (Fig. 2I) so that the fractional increments, Δ*r*, and fractional cell-to-cell heterogeneity are concisely presented. Quantification of the fractional cell-to-cell heterogeneity structure can be performed for any type of signaling data (see *Methods* and *SM*).

### Different types of PBMCs exhibit very similar logarithmic dose-responses

FRA overcomes the shortcomings of conventional analytical approaches to dose-response data by accounting for cell-to-cell heterogeneity and multivariate data. To determine the kinds of biological information that can be uncovered, we performed FRA for IFN-*α*2a dose-responses in specific types of PBMCs, assuming that all five measured pSTATs jointly constitute a cell’s response. The FRC and fractional cell-to-cell heterogeneity (Fig. 3A,B) are very similar for all cell types. Counter to what might be expected intuitively, and despite the differences seen in the conventional analysis (Fig. 1A-D), the dose responses in different cell types follow the same logarithmic pattern, identifying a phenomenon that governs the behavior of cellular populations in our system.

**Figure 3.**
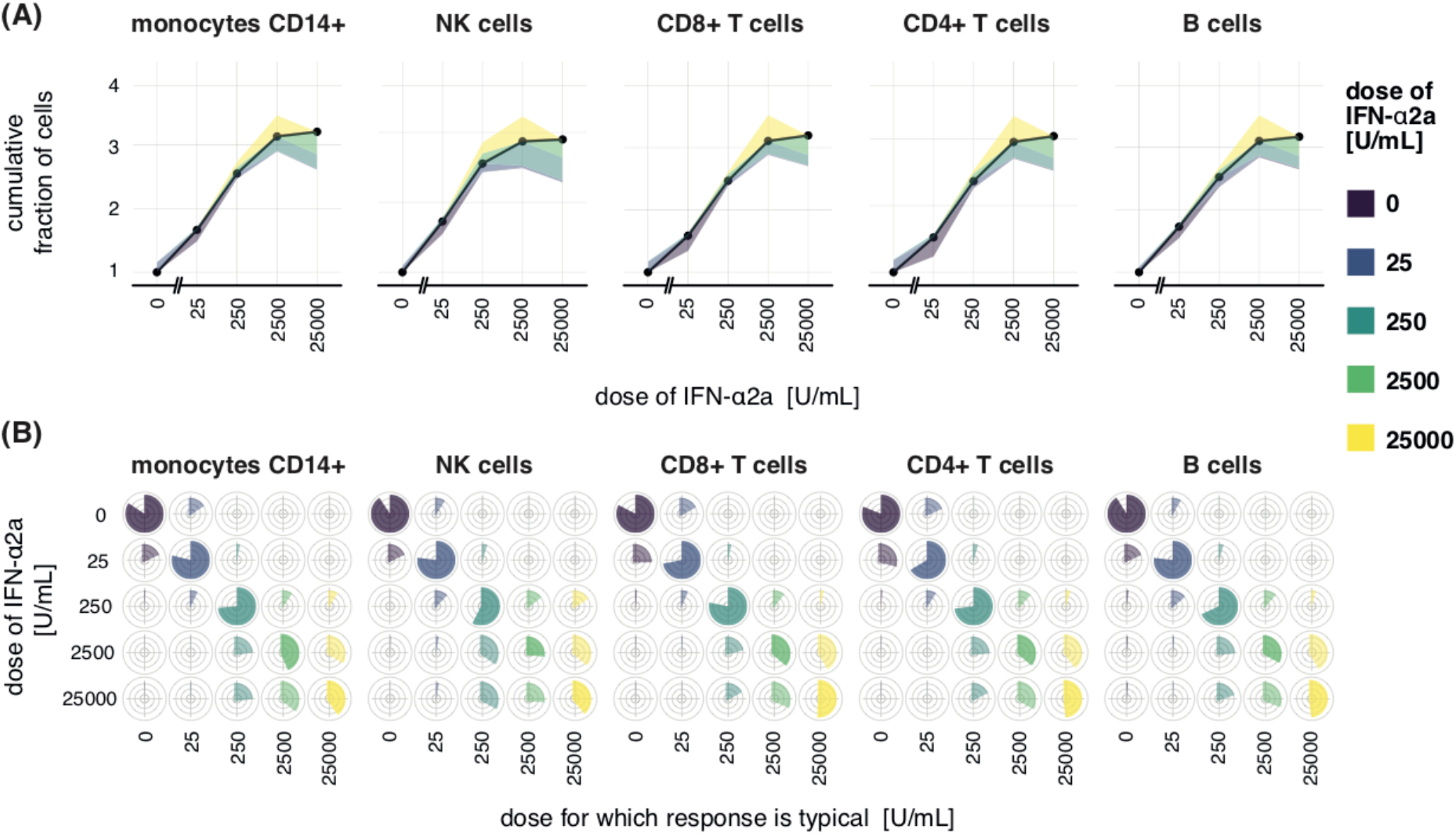
Different types of PBMCs exhibit highly similar dose-responses to IFN-*α*2a. **(A)** FRA of IFN-α2a responses. Here, levels of all pSTATs were assumed to jointly constitute cell’s response. Fig. S5 shows FRA for individual STATs. **(B)** Pie-charts of the cell-to-cell heterogeneity structure used to plot color bands in (A). Cell-to-cell heterogeneity is shown as pie-charts, in addition to (A), in order to clearly visualize similarity between the cell types.

For all cell types the FRC is linear and increases at the same rate with respect to the log of the dose, which means that the fraction of cells showing different responses is proportional to the dose fold-change, over a broad range of doses, i.e., from 0–2500 U/mL. The linear increase of the FRC demonstrates that the fraction of cells that exhibit different responses is very similar from 0–25 U/mL, from 25–250 U/mL, and from 250–2500 U/mL. For each subsequent dose change, Δ*r* ≈ 0.5 so that 50% of cells have different responses. A given fold change in the dose induces a different response in the fixed fraction of cells, across a broad range of doses. Therefore, cellular populations are sensitive to fold-changes in the dose as opposed to additive changes.

Formally, FRC scales as the log of the dose

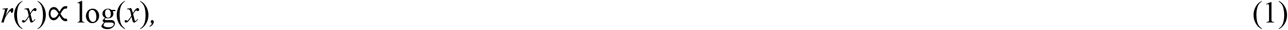

which given incremental approximation, Δlog(*x*) = log(*x*+Δ*x*) − log(*x*) ≈ Δ*x/x*, implies fold-change sensitivity in the population

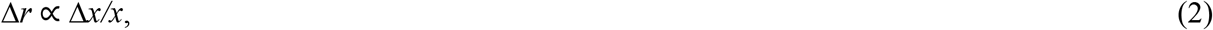

which in the studied system universally describes dose-responses in populations of different cell types. The FRA, therefore, condenses the description of the complex system into a simple quantitative formula. Furthermore, FRA uncovered that the number of programmed response distributions, i.e. maximal value of FRC, and the fractional cell-to-cell heterogeneity structure are very similar for all cell types. This similarity indicates that the immune system may precisely control responses of fractions of cells rather than responses of individual cells. In multicellular organisms, a fraction of cells with a given response level is a biologically essential response variable. For example, the outcome of a viral infection in a tissue depends on the number of NK cells with given response levels and induced cytotoxic activity. Our analysis revealed that in the studied system the fraction of cells that have responses in a specific range is not only tightly controlled in the population of a given cell type but is controlled in the same way across different cell types, as opposed to responses of individual cells that are largely heterogeneous within one cell type and across cell types. The role for controlling the fractions of cells with specific responses can, in principle, be tested by perturbing cell-to-cell heterogeneity through genetic manipulation and observing the phenotypic effects on the performance of the immune system.

### Logarithmic dose responses are a general property of cytokine signaling

To explore how generally applicable FRA is, we examined responses to cytokines IFN-*γ*, IL-10, and TNF-*α*^4,27^. As IFN-*γ* and IL-10 are implicated in macrophage phenotypic diversity^32^, we used the human monocyte cell line U937, differentiated into macrophage-like cells, and immunostaining to measure responses via nuclear levels of the key signaling effectors, phosphorylated STAT1 for IFN-*γ*, and phosphorylated STAT3 for IL-10 at 30 minutes after stimulation (Fig. 4A-B). Live confocal imaging and a murine embryonic fibroblasts cell line stably expressing fluorescent NF-*κ*B complex^4^, a key TNF-*α* signaling effector, were used to measure TNF-*α* responses in individual cells over time (Fig. 4C). For IFN-*γ*, distributions of responses shifted gradually towards higher values as the dose increases, referred to as the graded response^33–35^. For IL-10 the distributions flattened over a broad region as the dose increases, reflecting the higher number of responding cells for high doses, with the dose having a limited impact on the level of the response, similar to a binary system^2,35^ where responses aggregate in “on” and “off” regions. On the other hand, TNF-*α* responses are given as time-series, therefore, their characteristics cannot be directly observed.

**Figure 4.**
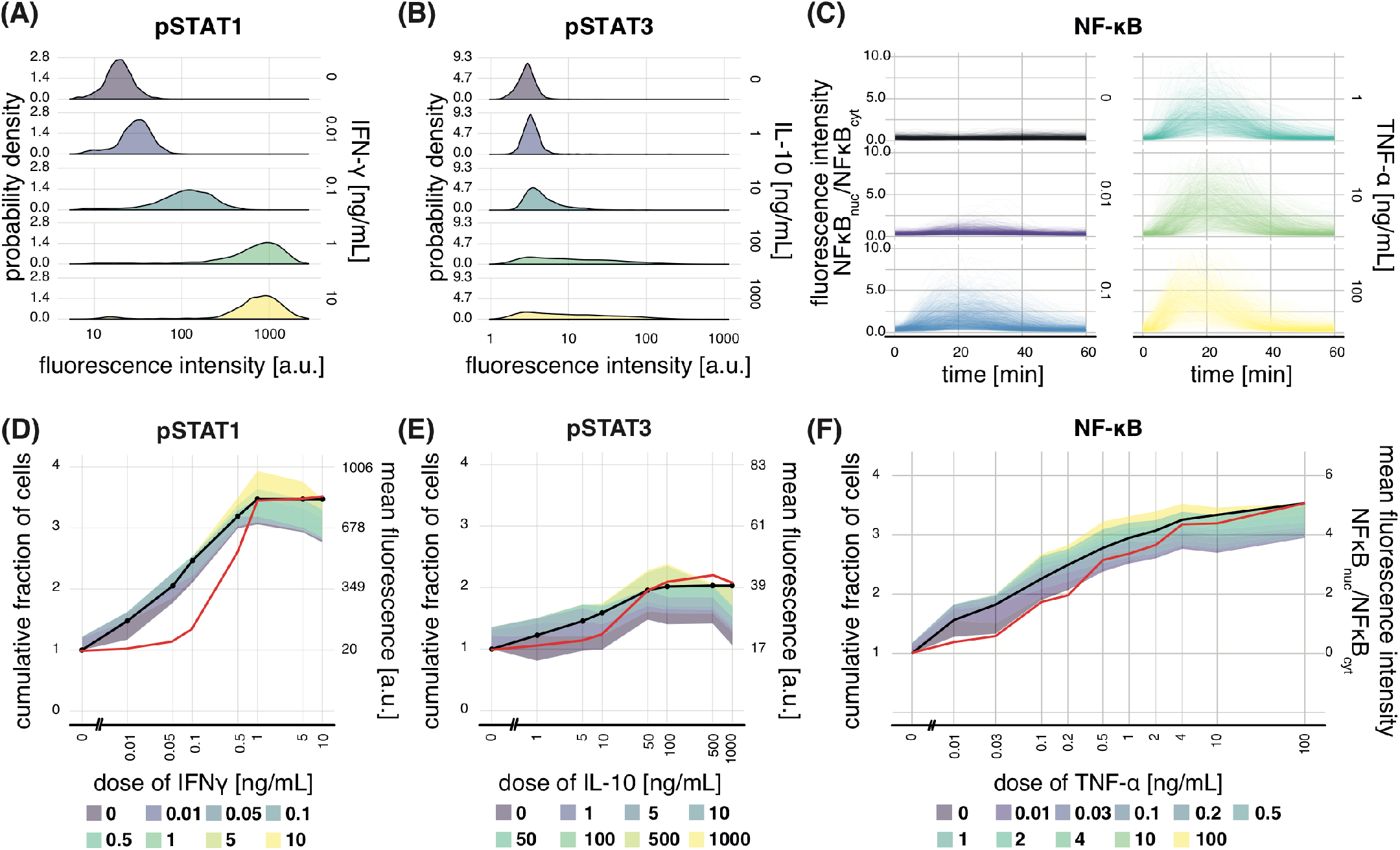
IFN-*γ*, IL-10, and TNF-*α* exhibit logarithmic dose responses in cell lines. **(A)** Distributions of responses 30 min after stimulation with different doses of IFN-*γ* in terms of nuclear pSTAT1 as measured with confocal microscopy imaging and immunostaining in the U937 cell line. Responses are expressed as mean fluorescence of nuclear pixels. Selected doses are shown. See Fig. S6 for all doses. **(B)** The same as in (A) but for pSTAT3 after stimulation with IL-10. **(C)** Temporally resolved responses of individual murine embryonic fibroblasts to increasing concentrations of TNF-*α*. Each line corresponds to a single cell. Responses are expressed as the ratio of the nuclear-pixels fluorescence average to the cytoplasmic-pixels fluorescence average. Selected doses are shown in the panel. See Fig. S7 for all doses. Measurements were taken every 3 min in a murine embryonic fibroblast cell line stably expressing the p65 protein component of the NF-*κ*B complex fused with the fluorescent dsRed protein. **(D)** FRA of IFN-*γ* responses. For comparison, the red line presents the mean response with y-axis on the right. The pie-chart of the plotted cell-to-cell heterogeneity structure is shown in Fig. S8. **(E)** Same as in (D) but for IL-10 responses. **(F)** FRA of the temporally resolved responses to TNF-*α*. The red line presents the mean response at the time of maximal response, i.e., at 18 min.

Despite qualitative differences in the responses, and type of data used, FRC increases nearly linearly with respect to the log of the dose for all three cytokines. Therefore, similarly to IFN-*α*2a in PBMCs, IFN-*γ*, IL-10, and TNF-*α* responses are sensitive to fold-changes in the dose, as opposed to additive changes, suggesting that this mode of response may be a more universal biological pattern that describes cytokine signaling in cellular populations. The qualitative differences in the responses to IFN-*γ* and IL-10 cytokines are reflected in a higher rate of increase of the FRC and narrower bands around FRC for IFN-*γ* compared to IL-10. The faster increase of the FRC for IFN-*γ* reflects a higher fraction of cells that exhibit different responses as the dose increases (Fig. 4A,B). The broader bands for IL-10 correspond to the bigger overlaps between the response distributions corresponding to different doses. For TNF-*α*, the rate of the increase of FRC as well as width of the bands around FRC are more similar to IFN-*γ* than for IL-10, which could not be determined directly from time-series data. Further, the different maximal values of the FRC for different cytokines revealed that signaling systems differ in terms of the number of programmed response distributions. These properties of dose-response are not captured by mean responses plotted for comparison as red line (Fig. 4D-F).

## Discussion

Sensitivity to dose fold-changes in populations of cells resembles the empirical Weber-Fechner law that characterizes the performance of many psycho-physiological sensory systems. Minimal detectable stimulus change, Δ*x*, in the sense of weight, hearing, vision, and smell, has been observed to be of fold-type. Our analysis shows, therefore, that cellular populations follow the same pattern, originally observed at the macro-scale in experimental psychology and neuroscience, and also more recently in conventional analysis of some cellular signaling systems, including bacterial chemotaxis and TGF-β signaling^12,36–38^. Therefore, the way cellular populations respond to stimuli is quantitatively similar to the way we perceive differences in certain sensations (weight/light). Weber-Fechner law is a pattern that can arise from a range of different mechanisms^39,40^ with the underlying neural implementations still being discovered^40,41^. Here also, a mechanistic explanation of the fold-change sensitivity of cellular populations is not clear and remains to be determined.

Overall, FRA delivers a concise representation of complex single-cell data, which is particularly relevant for high-throughput techniques, which are increasingly allowing the measurement of a high number of parameters per cell, generating vary large, high-dimensional datasets^42^. The high information content of multivariate, single-cell measurements makes biological discoveries more likely. On the other hand, however, insights may be difficult to extract due to data complexity. Therefore, making use of the increasing amount of single-cell high-throughput measurements requires approaches that can extract relevant insights in spite of complexity. FRA, not being limited to cytokine signaling, proteomic data or dose responses, enables the systematic investigation of single-cell high-throughput data in a wide range of situations, in which responses are measured in single-cells at any “-omics” scale. Therefore, FRA should yield insights into the structure of signaling heterogeneity in immunology, developmental biology, cancer research, and diverse other fields in which response analysis in single-cells is of relevance.

## Methods

### Software implementation

The methodology to perform and visualize FRA is provided as a user-friendly R-package available for download at http://github.com/sysbiosig/FRA. The package contains an installation guide and a brief user manual.

### Formal definition of the FRC

Consider a series of doses *x*_1_,…,*x*_*i*_,…,*x*_*m*_ and denote a single cell response as *y*. Depending on the context, *y*, may be a number or a vector, e.g. the level of one or more measured signaling effectors. Suppose that responses to a given dose, *x*_*i*_, are represented as the probability distribution,

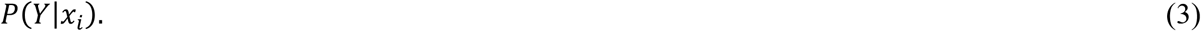

The FRC is then formally defined as

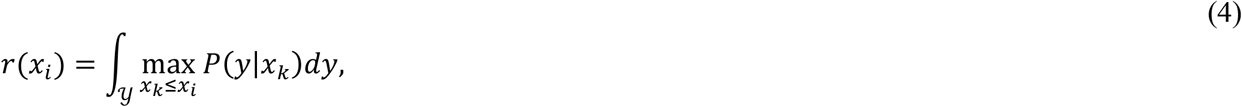

where integration takes place over 𝒴, the set of all possible responses, *y*. The integral quantifies the area under the curve (or under surface for multivariate data), with respect to *y*, defined as 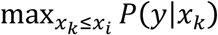. For the calculations shown in Fig. 2 the integration corresponds to the calculation of the area of the gray regions in panels D-E. As explained in the *SM*, the FRC defined as above is closely related Rényi min-information capacity.

### Formal definition of typical fractions

Having the responses represented in terms of the probability distribution, Eq. 3, we can define which responses, *y*, are typical to any of the doses. Precisely, we define the response, *y*, to be typical for dose *x*_*j*_ if it is most likely to arise for this dose, which writes as

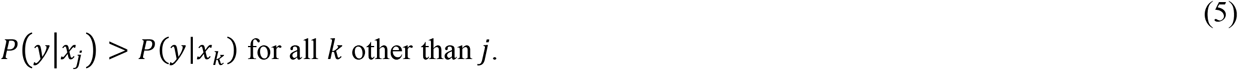

The above condition allows assigning any response, *y*, to a dose for which it is typical. Therefore, for a given dose, *x*_*i*_, we can identify what fraction of cells stimulated with this dose exhibits responses typical to any dose, *x*_*j*_, for *j* from 1 to *m*. These fractions, denoted as *v*_*ij*_, can be practically computed as explained below.

### Calculation of typical fractions

The fractions of cells stimulated with dose *i* that have responses typical to dose *j, v*_*ij*_, can be easily calculated from data regardless of the number of doses and the type of experimental measurements. We have that

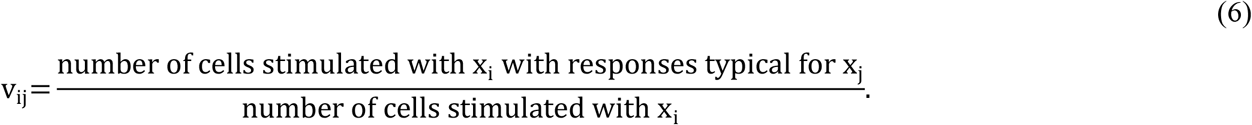

Calculation of typical fractions, *v*_*ij*_, with the above formula requires the possibility to examine the condition *P*(*y*|*x*_*j*_) *> P*(*y*|*x*_*k*_) for any experimentally observed response, *y*. The distributions *P*(*y*|*x*_*j*_) can be reconstructed from data using a variety of probability density estimators^43^. The use of the available estimators, however, might be problematic for multivariate responses^24,44^. We therefore propose a more convenient strategy. We replace the condition *P*(*y*|*x*_*j*_) *> P*(*y*|*x*_*k*_) with an equivalent condition that is computationally much simpler to evaluate. Precisely, we propose to use the Bayes formula

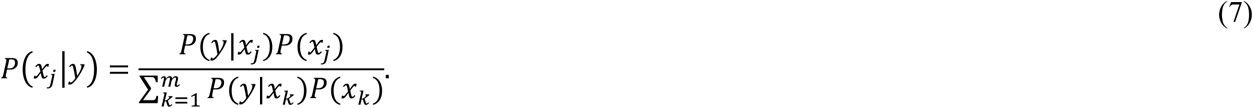

If we set the equiprobable prior distribution, i.e., *P*(*x*_*j*_)= 1*/m*, we have that *P*(*y*|*x*_*j*_) is proportional to *P*(*x*_*j*_|*y*) and the condition *P*(*y*|*x*_*j*_) *> P*(*y*|*x*_*k*_) is equivalent to

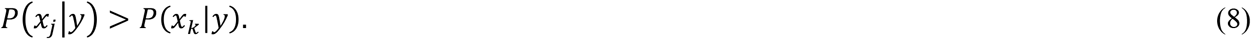

The above strategy allows avoiding estimation of the response distributions, *P*(*y*|*x*_*j*_), from data. For continuous and multivariate variable *y* the estimation of *P*(*x*_*j*_|*y*) is generally simpler than estimation of *P*(*y*|*x*_*j*_)^24,43^. Precisely, an estimator 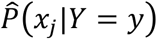 of the distribution *P*(*x*_*j*_|*y*) can be built using a variety of Bayesian statistical learning methods. For simplicity and efficiency, here we propose to use logistic regression, which is known to work well in a range of applications^43^. In principle, however, other classifiers could also be considered. The logistic regression estimators of *P*(*x*_*j*_|*Y* = *y*) arise from a simplifying assumption that log-ratio of probabilities, *P*(*x*_*j*_|*Y* = *y*) and *P*(*x*_*m*_|*Y* = *y*) is linear. Precisely,

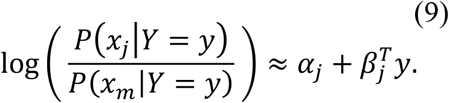

The above formulation allows fitting the logistic regression equations to experimental data, i.e., finding values of the parameters, *α*_*j*_ and *β*_*j*_ that best represent the data. The fitted logistic regression model allows assigning cellular responses to typical doses based on condition given by Eq. 8. Formally, the fractions *v*_*ij*_ defined by Eq. 6 are calculated as

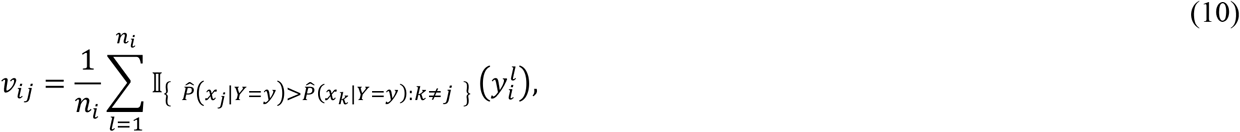

where *n*_*i*_ is the number of cells measured for the dose *x*_*i*_, *y*^*l*^_*i*_ denotes response of the *l*-th cell, and 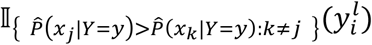 is equal 1 if 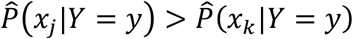 for any *k* ≠ *j* and 0 otherwise.

### Computation of the FRC

Calculation of the SCRC can be conveniently performed using the typical fractions, as defined above, rather than through integration of Eq. 4. Precisely, to calculate the FRC for the dose, *x*_*i*_, consider doses *x*_1_,…,*x*_*i*_ in isolation from higher doses. Then, the sum of typical fractions *v*_11_,…,*v*_*ii*_ is equivalent to FRC for the dose *x*_*i*_

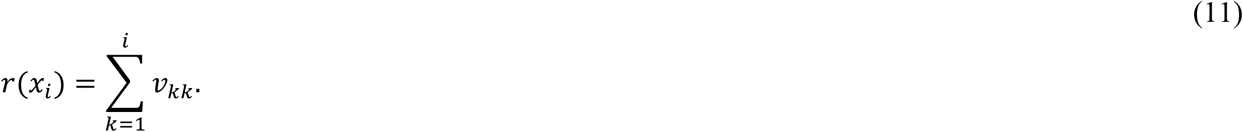

The equivalency of the above equation and Eq. 4 is explained in the *SM*.

### Mass cytometry (CyTOF)

PBMCs were isolated from the peripheral blood of healthy adult donors using Lymphoprep (Stemcell Technologies), according to the manufacturer’s instructions. Cells were washed in serum-free RPMI then resuspended at 10^7^ cells/mL in serum-free RPMI containing 0.5 mM Cell-ID Cisplatin (Fluidigm) and incubated at 37°C for 5 min. Cells were washed with RPMI containing 10% (v/v) FCS (Sigma) and 2 mM L-Glutamine (R10), centrifuging at 300 x g for 5 min before being resuspended to 6 x 10^7^ cells/mL in R10 and rested at 37^°^C for 15 min. 50 mL of cells (3 x 10^6^ cells) were transferred to 15 mL falcon tubes for stimulation and antibody staining. Antibodies are listed in Supplementary Table 1. Staining for CD14, CCR6, CD56, CD45RO, CD27, CCR7, CCR4 and CXCR3 was done before stimulation/fixation for 30 min in R10 at 37^°^C. Cells were stimulated with 0, 25, 250, 2500 or 25000 U/mL recombinant human IFN-*α*2a (PBL Assay Science, #11100-1) diluted in R10 for 15 min at 37^°^C. After washing with 5 mL cold Maxpar PBS (Fluidigm), cells were fixed with 1 X Maxpar Fix I Buffer (Fluidigm) for 10 min at RT before being washed with 1.5 mL Maxpar Cell Staining Buffer (CSB, Fluidigm). All centrifugation steps after this point were at 800 x g for 5 min. Cells were barcoded using Cell-ID 20-Plex Pd Barcoding Kit (Fluidigm), according to the manufacturer’s instructions, and washed twice with CSB before samples were pooled and counted. All further steps were performed on the pooled cells. Fc receptors were blocked using Fc Receptor Binding Inhibitor Antibody (eBioscience, #14-9161-73) diluted 1:10 in CSB for 10 min at RT. Surface antibody staining mixture was added directly to the blocking solution and incubated for 30 min at RT. Cells were washed twice with CSB, resuspended in ice-cold methanol and stored at -80^°^C overnight. After washing twice with CSB, cells were stained with intracellular antibody staining mixture for 30 min at RT before two further washes in CSB. Cells were resuspended in 1.6% (v/v) formaldehyde (Pierce, #28906) diluted in Maxpar PBS and incubated for 10 min at RT. Cells were resuspended in 125 mM Cell-ID Intercalator (Fluidigm) diluted in Maxpar Fix and Perm Buffer (Fluidigm) and incubated overnight at 4^°^C. Compensation beads (OneComp eBeads Compensation Beads, Invitrogen, #01-1111-42) stained with 1 mL of each antibody were also prepared. The next day, cells and compensation beads were washed twice with CSB and twice with Maxpar water (Fluidigm), mixed with a 1:10 volume EQ Four Element Calibration Beads (Fluidigm) before acquisition on a Helios Mass Cytometer (Fluidigm) using the HT injector. Data were normalized, randomized and concatenated using CyTOF Software v6.7 (Fluidigm). Compensation and de-barcoding was performed using the CATALYST package^45^. Single, live, CD45+ cells were gated using Cytobank (Cytobank, Inc.)

### U937 cells

U937 cells (CRL-1593.2, ATCC), a human monocyte cell line, were cultured under standard conditions at 37°C in a humidified atmosphere of 5% CO_2_/95% air in low glucose Roswell Park Memorial Institute 1640 (RPMI 1640, Corning, #10-040-CV) medium supplemented with 10% fetal bovine serum (FBS, ThermoFisher, #10500064) and 1% penicillin-streptomycin solution (P/S, ThermoFisher, #15140122). For macrophage differentiation U937 cells were suspended in medium with 20 ng/mL phorbol 12-myristate 13-acetate (PMA, Sigma Aldrich, #P1585) and plated in 96-well microplates with *µ*Clear®flat bottom (Greiner, #655090) in density 2×10^4^ cells per well. After 24 h medium with PMA was removed and fresh medium was added to cells. 72 h after seeding on 96-well microplates differentiated cells were incubated with recombinant human interferon gamma (IFN-*γ*, ThermoFisher, #PHC4031) at concentrations 0-10 ng/mL or recombinant human interleukin 10 (IL-10, PeproTech, #200-10) at concentrations 0-1000 ng/mL for 30 min. Afterwards, cells were fixed with 3.7% paraformaldehyde (PFA, Sigma Aldrich, #P6148) for 10 min in room temperature, RT, then permeabilized with 90% ice-cold methanol (Sigma, #322415), for 30 min in -20°C, blocked with 5% bovine serum albumin (BSA, Merck, #821006) and 0.3% Triton X-100 (Sigma Aldrich, #T9284) for 1 h in RT, and incubated with primary antibody - phospho-STAT1 (Tyr701) (pSTAT1, Cell Signaling, #9167) diluted 1:100 or phospho-STAT3 (Tyr705) (pSTAT3, Cell Signaling, #4113) diluted 1:200 in 1% BSA with 0.3% Triton X-100 for 18 h in 4°C. Next day, cells were incubated with an appropriate secondary antibody - Alexa Fluor 488 (Life Technologies, #A-21206) or Alexa Fluor 555 (Life Technologies, #A-31570) diluted 1:500 in 1% BSA with 0.3% Triton X-100 for 1.5 h in RT and stained with 2 *µ*g/mL 4’,6-diamidino-2-phenylindole (DAPI, Sigma Aldrich, #D9542) for 10 min in RT. The fluorescence signal was imaged and quantified using automated confocal microscope (Pathway 435, BD) and analyzed with CellProfiler and ImageJ.

### Murine immortalized fibroblasts

Murine embryonic fibroblasts cell line, NIH/3T3 (CRL-1658, ATCC), expressing fluorescent fusion proteins relA-dsRed as wells H2B-GFP for nuclei identification were cultured in incubator under standard conditions at 37°C in a humidified atmosphere of 5% CO2/95% air. The cell line was kindly provided by prof. S. Tay and was previously used in several studies, including^4,24^. The cells were cultured in high glucose Dulbecco’s Modified Eagle’s Medium without phenol red (DMEM, ThermoFisher, #21063029) supplemented with 10% fetal bovine serum (FBS, ThermoFisher, #10500064) and 1% penicillin-streptomycin solution (P/S, ThermoFisher, 15140122). Approximately 1.3 x 10^5^ cells were plated on 35-mm confocal dish for imaging. After 48 h in the incubator, cells were transferred to environmental chamber in a microscope. At time 0 medium was removed from cells and recombinant mouse tumor necrosis factor alpha (TNF-*α*, Sigma-Aldrich, #T7539) was added at concentrations 0-100 ng/mL as 5-minute pulse. Live imaging was performed using a confocal microscope, Leica TCS SP5 X. During single experiment images have been captured every 3 minutes over 1 hour in two channels simultaneously at 9 different positions on the plate. Experiment has been repeated at least four times to test reproducibility and to allow for a sufficient number of observations. Nuclear and cytoplasmic fluorescence (pixel mean) was then quantified from microscopic images. The response of each cell was then represented as the ratio of nuclear to cytoplasmic fluorescence in order to ensure robustness of measurements to changes in confocal plane over time. The data set is described in detail in^24^, where it was initially published.

## Acknowledgements

KN, JW, KEZ and MK were supported by Foundation for Polish Science within the First TEAM (First TEAM/2017-3/21) program co-financed by the European Union under the European Regional Development Fund. RER and JR were funded by the UK Medical Research Council [MRC core funding of the MRC Human Immunology Unit]. We thank Guy Riddihough for helpful comments during the preparation of this manuscript. MK and JR also thank Matteo Iannacone, the organizer of an EMBO workshop on Immunology, Bergamo 2017, that had initiated our collaboration.

## Supplementary Materials Legends

**S1 Text**. Supplementary information text contains expanded description of theoretical methods.

**S1 Figure**. Time-course of responses to INF-*α*2a in PBMCs, corresponds to Fig. 1.

**S2 Figure**. Dose-responses to IFN-*α*2a in PBMCs presented as t-SNE plots, corresponds to Fig. 1B.

**S3 Figure**. Dose-responses to IFN-*α*2a in PBMCs shown as distributions of pSTATs, corresponds to Fig. 1D.

**S4 Figure**. FRCs counts the number of distinct response distributions, corresponds to Fig. 2.

**S5 Figure**. FRA of IFN-*α*2a responses plotted for individual STAT proteins, corresponds to Fig. 3.

**S6 Figure**. IFN-*γ*, IL-10 response distributions, corresponds to Fig. 4A,B.

**S7 Figure**. Temporally resolved responses to TNF-*α*, corresponds to Fig. 4C.

**S8 Figure**. Pie-charts of the cell-to-cell heterogeneity structure used to plot color bands in Fig. 4D-F.

**S1 Table**. Antibodies used for mass cytometry experiments.

## Supplementary text (S1 Text)

### Interpretation in terms of information-theory

The FRA is inspired by the mathematical theory of information, specifically the concepts of Rényi min-information^46–49^. Broadly speaking, one of the concerns of information theory is to quantify how much information about an input variable *X* is contained in an output variable *Y*. Typically, the input variable denotes signals (messages) that need to be decoded from the output variable^50–52^. To explain how the FRA relates to information theory, assume that doses, *x*_1_,…,*x*_*i*_, represent signals that need to be decoded from the cellular responses, *y*, by an external observer. If response distributions to each dose were completely distinct, each of the *i* doses could be decoded by the observer without error, as responses to each dose correspond to a different range. On the other hand, if distributions corresponding to different doses exhibited some overlap, a decoding strategy would be needed to translate a response back to the dose. It is well established in statistics, the best decoding strategy is the intuitive one: assign the signaling response to the dose for which it is most frequent^50^. Then, among cells stimulated with the dose *x*_*k*_, the faction that could be decoded correctly is the fraction that exhibits responses most frequent for *x*_*k*_, which in the paper is referred to as typical and quantified as *v*_*kk*_. It is shown in Fig. 2J for the three doses example. FRC, *r*(*x*_*i*_), as shown in Eq. 11 can be written as the sum of fractions of cells that can be decoded correctly

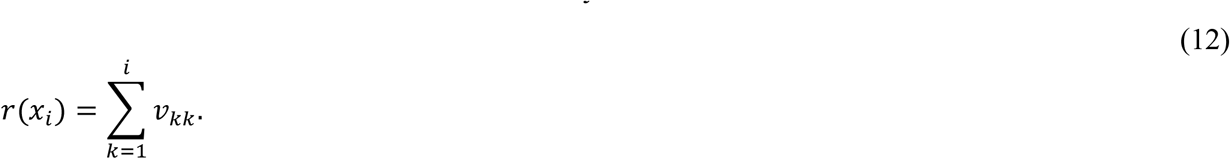

The fact that *r*(*x*_*i*_) is the sum of fractions that can be decoded correctly implies that 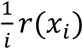 is the probability that an observer would guess correctly the dose of a randomly selected cell among cells stimulated with any of the *i* doses. Therefore, the product of the of number of doses, *i*, and the probability of correct decoding 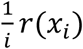, which equals *r*(*x*_*i*_), can be interpreted as the number of different doses that can be decoded correctly on average.

The above intuitive reasoning, which provided the interpretation of *r*(*x*_*i*_) in terms of the number of different doses that can be decoded correctly on average by the observer, is formalized within information-theory by the Rényi min-information. Rényi min-information, similarly to Shannon’s information, allows to quantify how much information about a variable *X* is transferred to a variable *Y*, Precisely, for a set of input signals *x*_1_,…,*x*_*i*_ and output responses distributed as *P*(*Y*|*x*_*i*_) the information transferred from *X* to *Y* is quantified by Rényi min-information capacity, 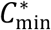, defined as^46,48,49^

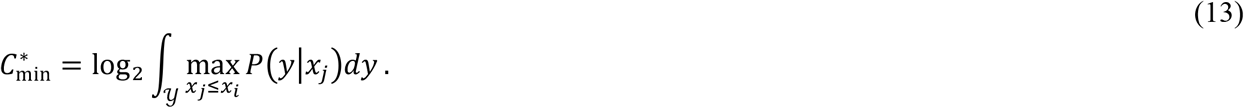

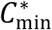 is expressed in bits of Rényi min-entropy^46,48,49,51^ and can be interpreted as the log_2_ of the number of different messages that can be transferred from *X* to *Y* in the sense of Rényi min-entropy.

We have, therefore, that

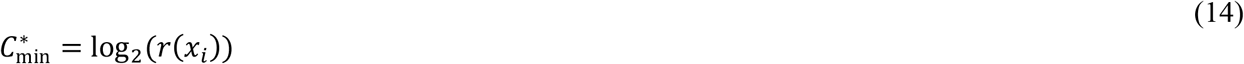

or equivalently

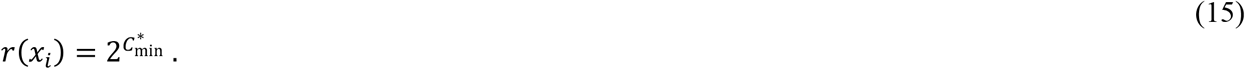

### Equivalency of Eq. 4 and Eq. 12

Formally, the fraction of cells stimulated with the dose *k* that can be decoded correctly is the fraction of the probability density, *P*(*y*|*x*_*k*_), with highest values among all doses *x*_1_,…,*x*_*i*_,

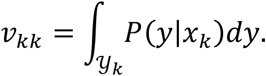

where 𝒴_*k*_ is the set of responses *y* typical for the dose k

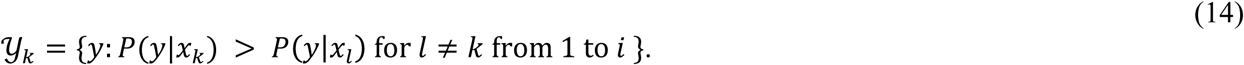

Therefore, the sum of fractions of cells that can be decoded correctly among doses *x*_1_,…,*x*_*i*_, gives the value of the FRC

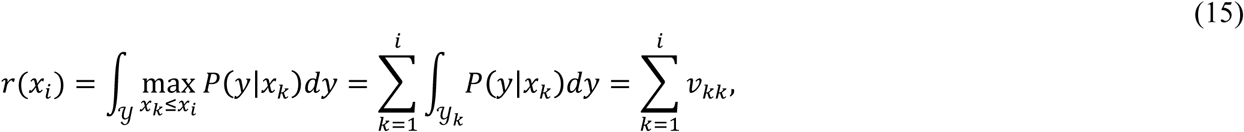

which demonstrates that definitions of FRC given by Eq. 4 and Eq. 12 are equivalent.

## Supplementary Figures

**Supplementary Figure 1.**
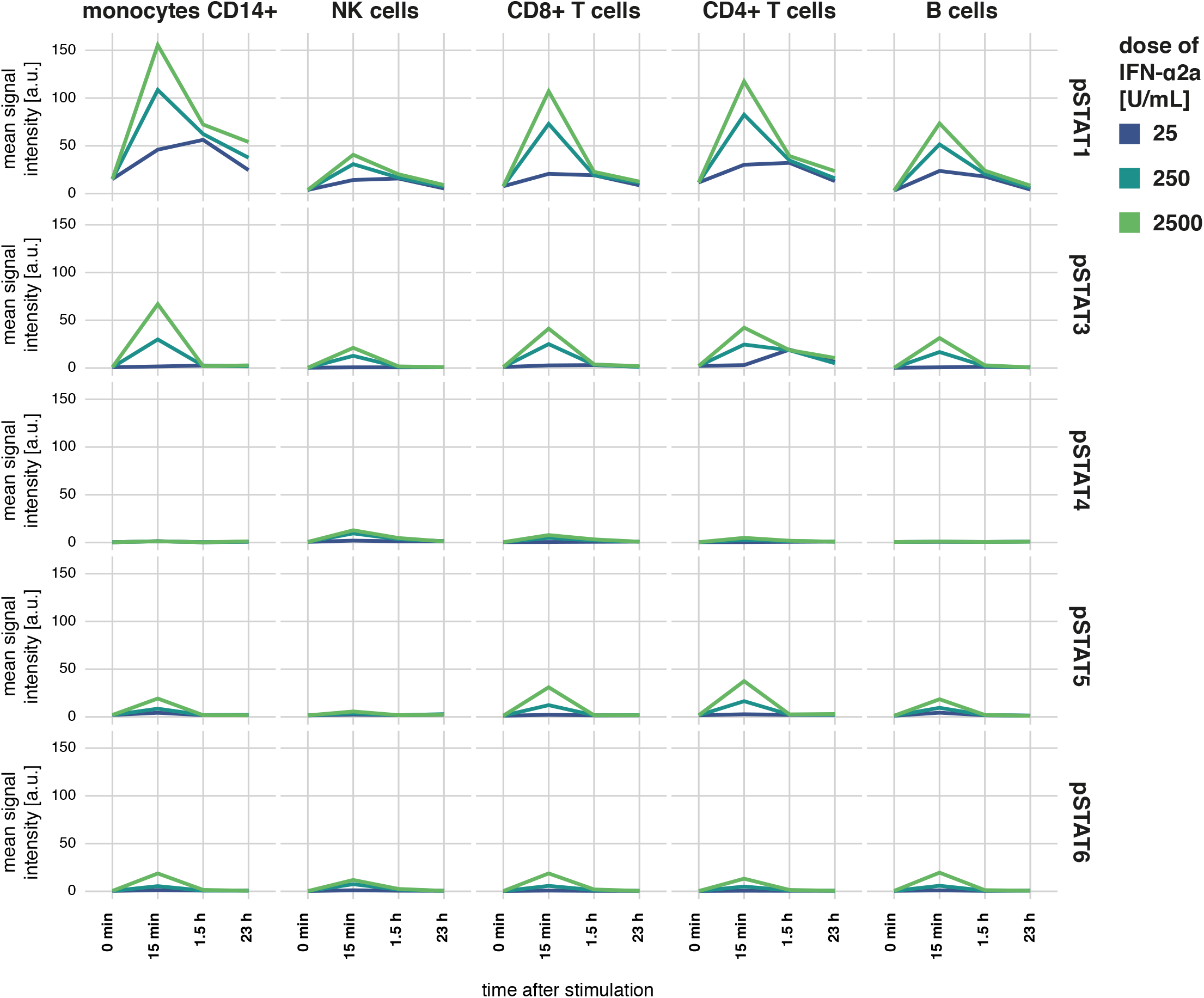
Time-course of responses to IFN-*α*2a in PBMCs. Population mean response (y-axis) of each cell type (columns) is shown in terms of whole cell levels of different pSTATs (rows), as measured with mass cytometry for different time points (x-axis) after stimulation with the indicted dose (color).

**Supplementary Figure 2.**
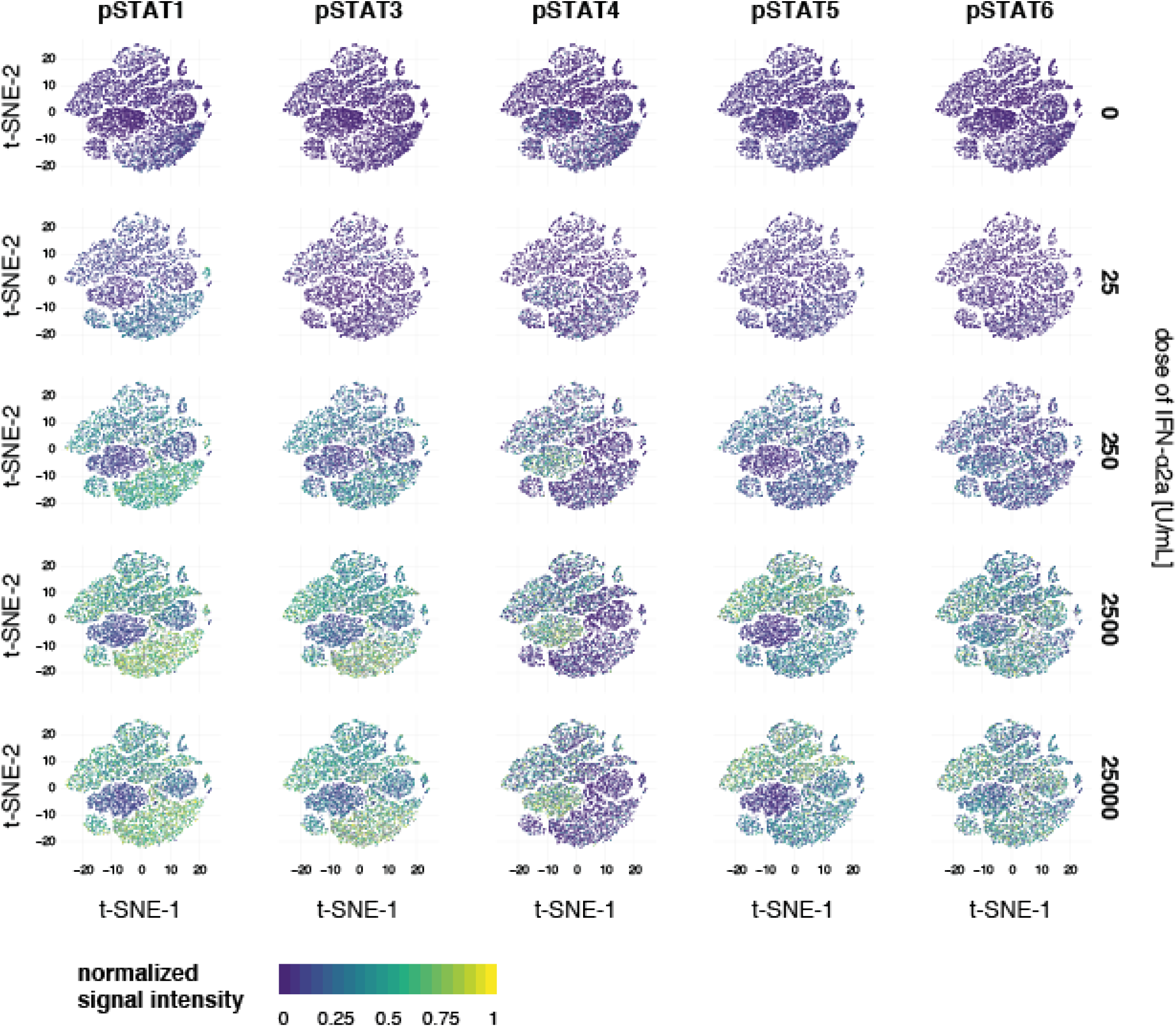
Dose-responses to IFN-*α*2a in PBMCs presented as t-SNE plots. Figure corresponds to Fig. 1B, where responses to selected doses are presented. Here all doses are shown.

**Supplementary Figure 3.**
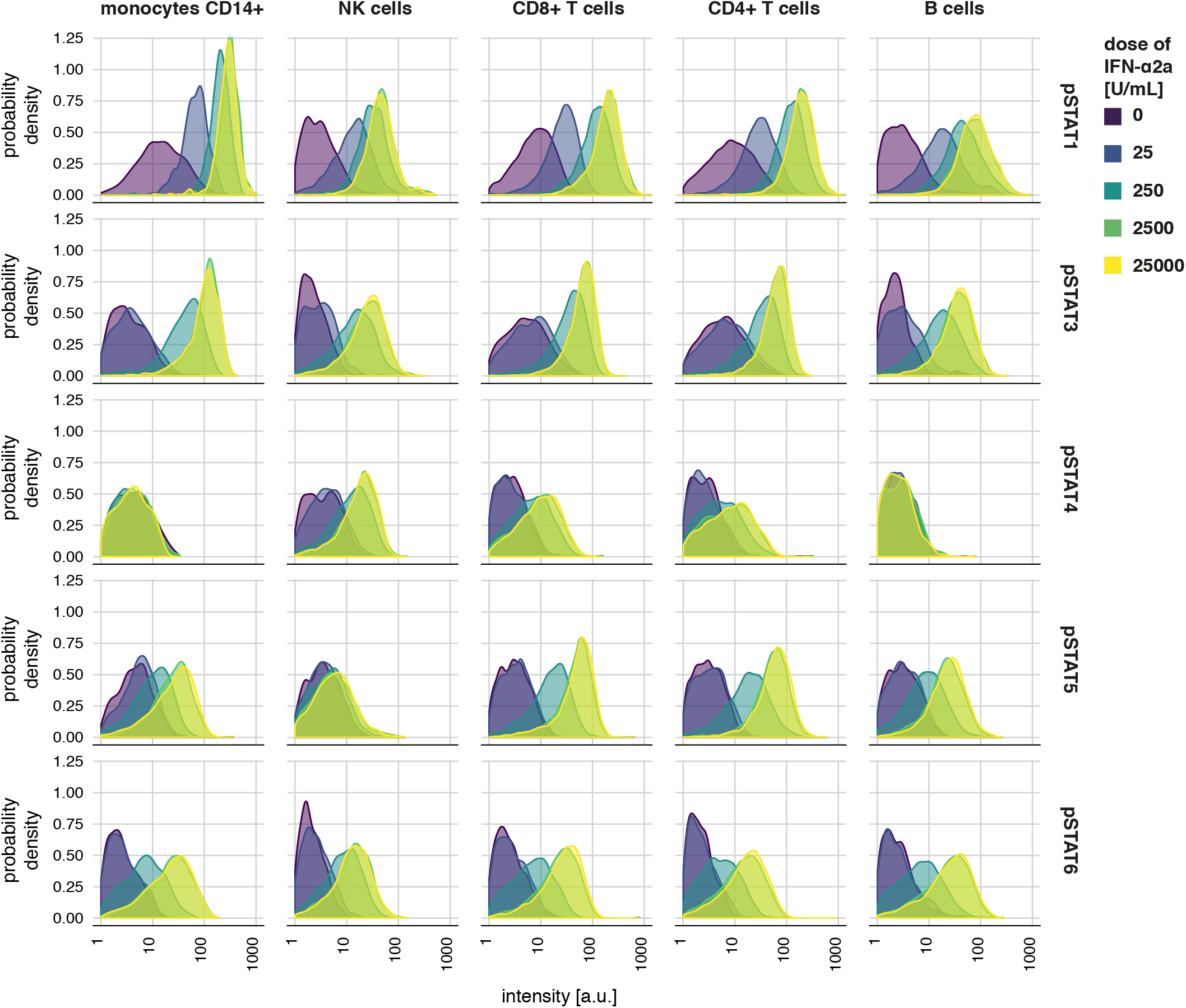
Dose-responses to IFN-*α*2a in PBMCs shown as distributions of pSTATs. Figure corresponds to Fig. 1D, where distributions of pSTAT1 and pSTAT5 are presented. Here all pSTATs are shown.

**Supplementary Figure 4.**
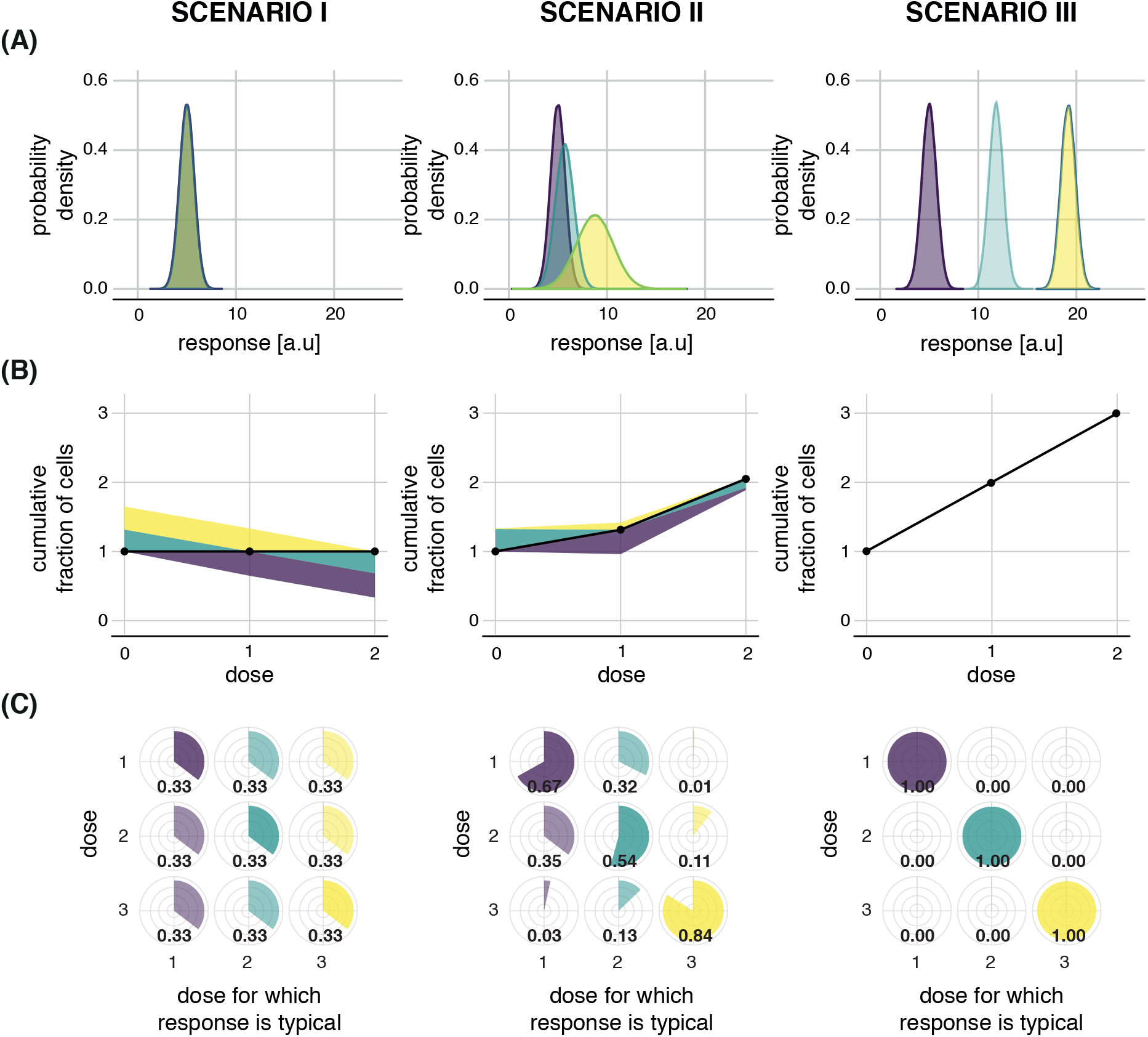
FRC counts the number of distinct response distributions. Figure corresponds to Fig. 2. **(A)** Distributions of responses in three hypothetical scenarios. Scenario I: completely overlapping responses; Scenario II: the same as in Fig. 2, i.e., partly overlapping responses; Scenario III: completely distinct responses. **(B)** Distributions of responses in three hypothetical scenarios. Scenario I: completely overlapping responses; Scenario II: the same as in Fig. 2, i.e., partly overlapping responses; Scenario III: completely distinct responses. **(C)** Pie-charts representing cell-to-cell heterogeneity structure in the three scenarios (columns).

**Supplementary Figure 5.**
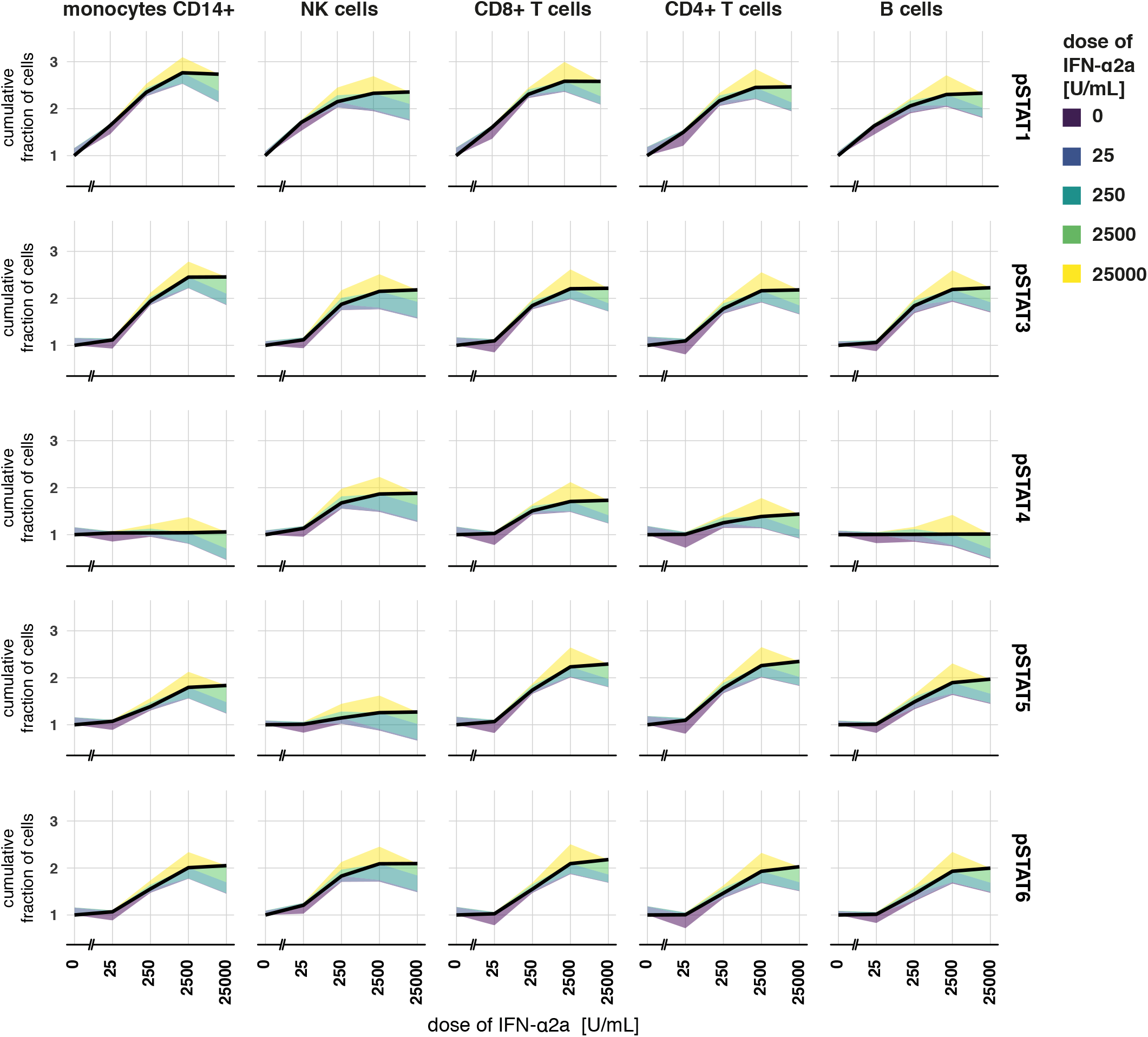
FRA of IFN-*α*2a responses for individual STAT proteins. Figure corresponds to Fig. 3A, where FRA were performed assuming that all pSTATs jointly constitute cell’s response. Here, each panel presents FRA for an individual pSTAT (rows) for different cell types (columns). Data used to plot each panel is shown in the corresponding panel of Supplementary Figure 3.

**Supplementary Figure 6.**
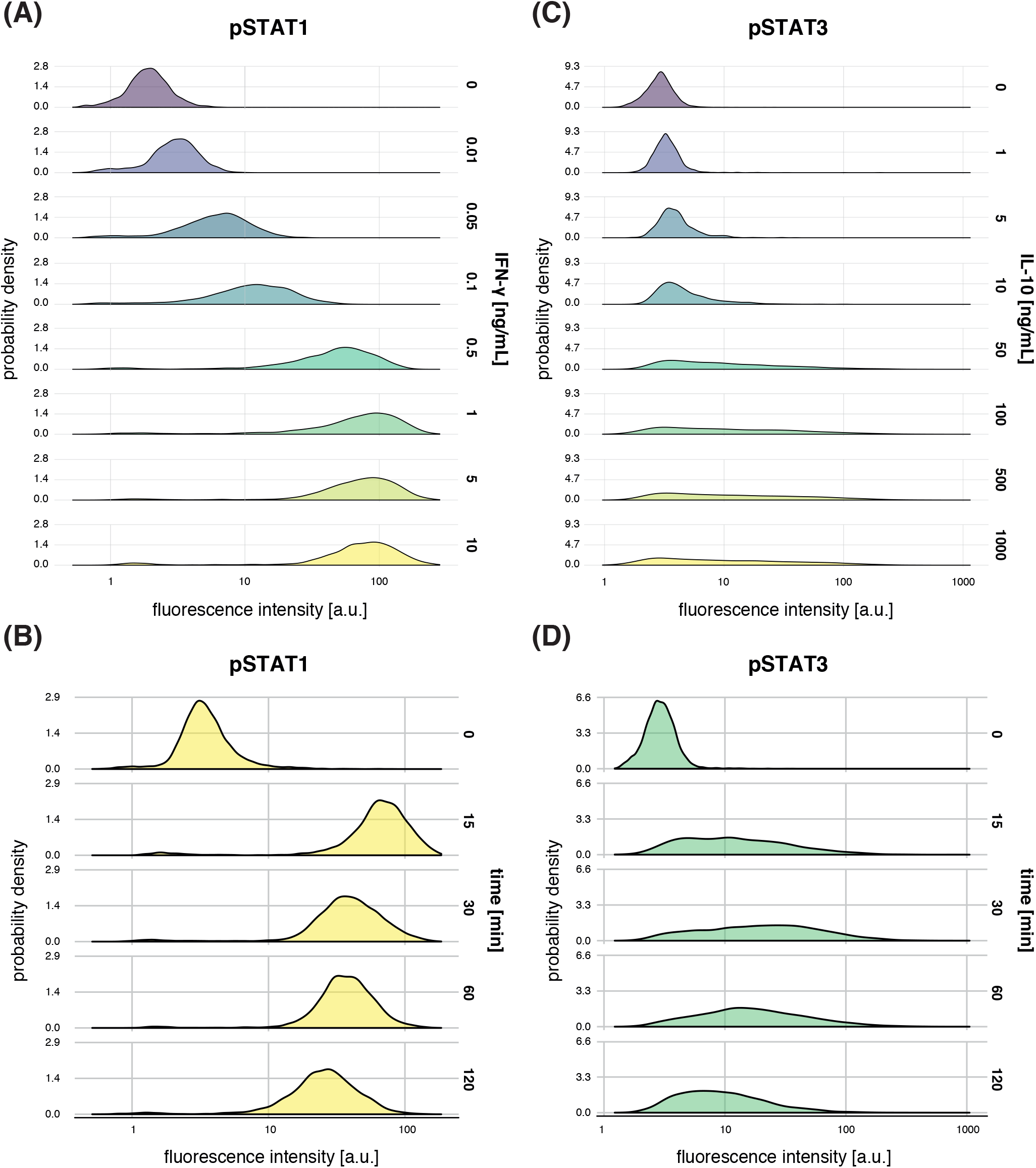
IFN-*γ*, IL-10 response distributions. Figure corresponds to Fig. 4A,B. **(A)** Response distributions to IFN-*γ* as in Fig. 4A were response distributions to selected doses are shown.Here all doses are shown. **(B)** Same as in (A) but for IL-10, corresponds to Fig. 4B. **(C)** Distributions of responses for different times after stimulation with 10 ng/mL of IFN-*γ*. **(D)** Distributions of responses for different times after stimulation with 100 ng/mL of IL-10.

**Supplementary Figure 7.**
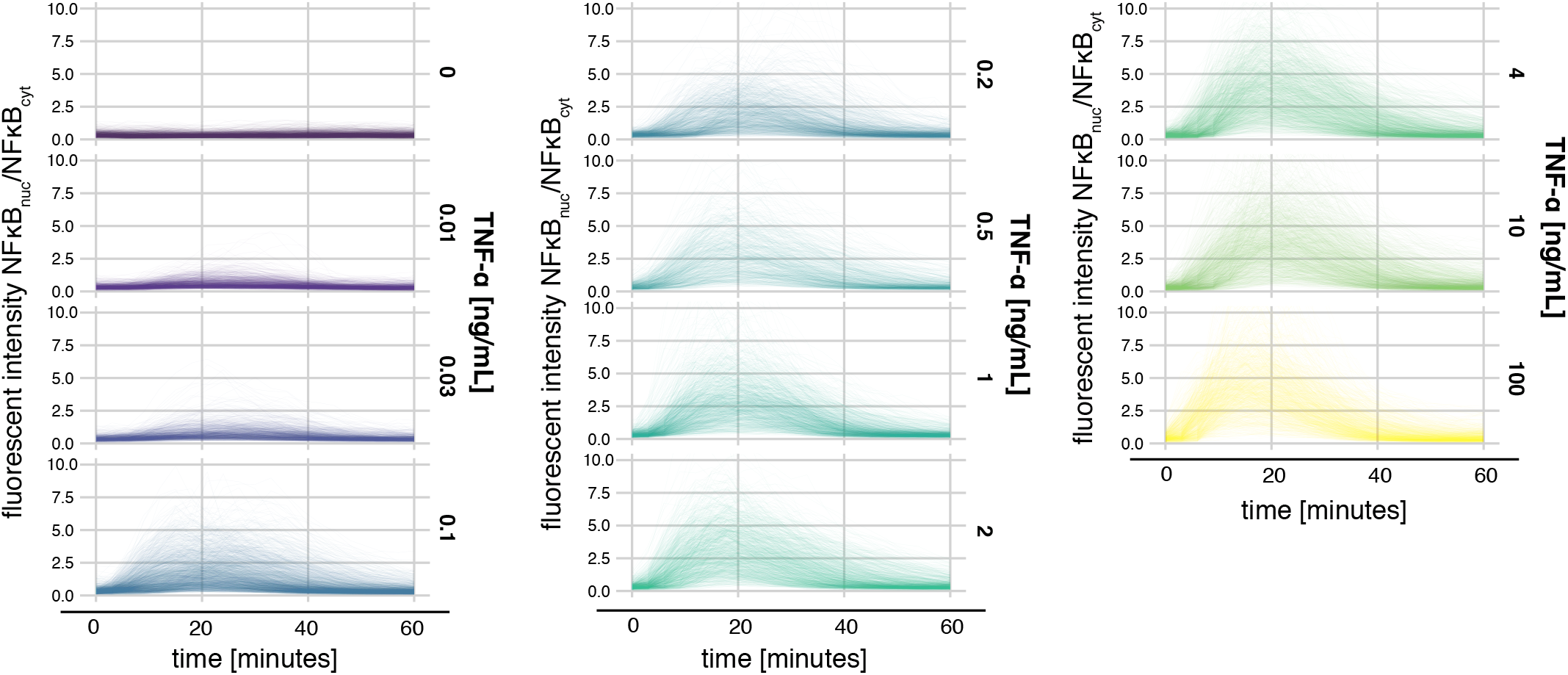
Temporally resolved responses to TNF-*α*. Figure corresponds to Fig. 4C, where responses to selected doses are presented. Here all doses are shown.

**Supplementary Figure 8.**
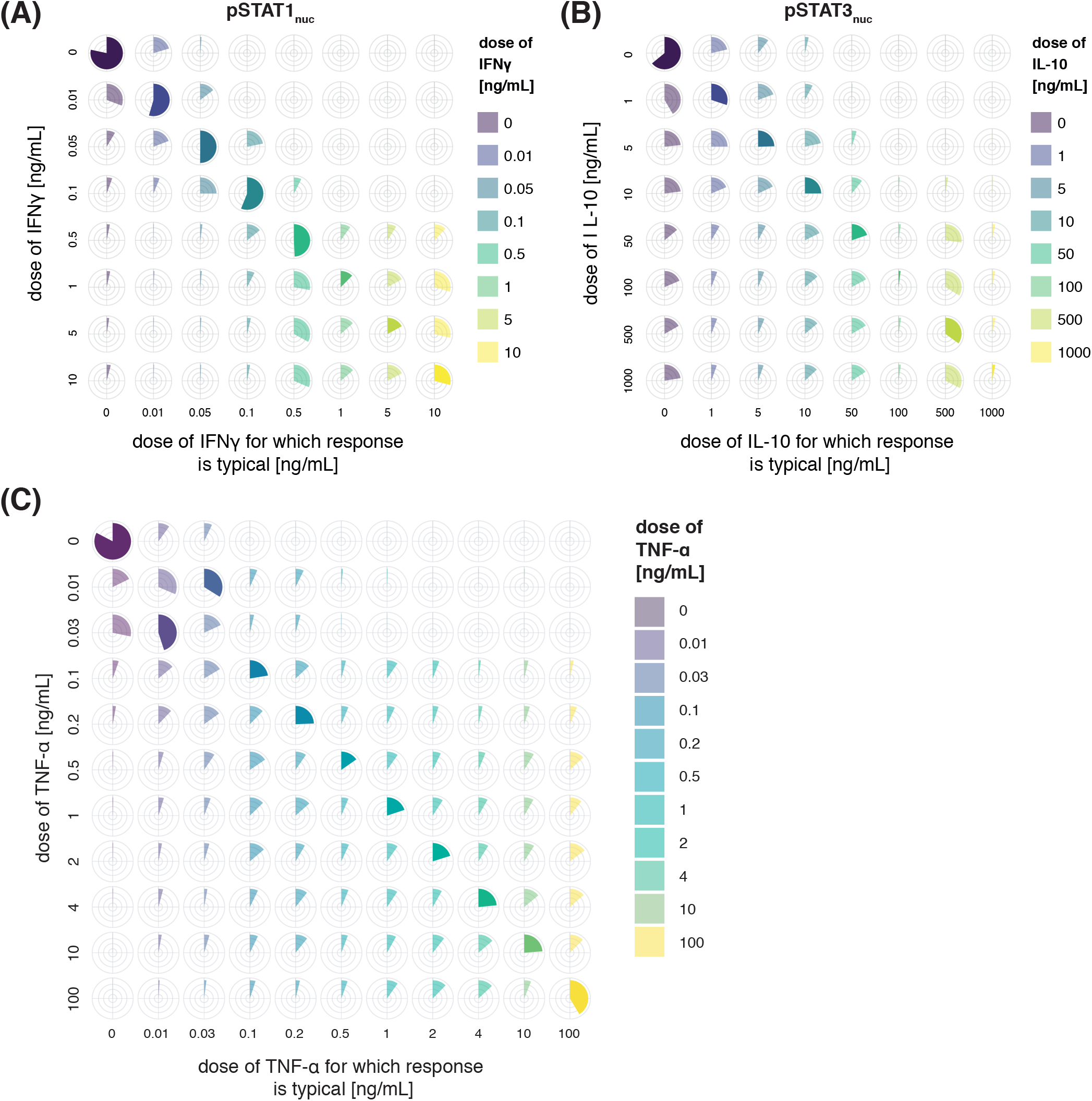
Pie-charts of the cell-to-cell heterogeneity structure used to plot color bands in Fig. 4D-F. **(A)** Cell-to-cell heterogeneity structure of IFN-*γ* responses, corresponds to Fig. 4D. **(B)** Cell-to-cell heterogeneity structure of IL-10 responses, corresponds to Fig. 4E. **(C)** Cell-to-cell heterogeneity structure of NF-*κ*B responses, corresponds to Fig. 4F.

## Supplementary Table

**Supplementary Table 1.** Antibodies used for mass cytometry experiments.

| Target | Clone | Metal label | Company | Catalogue number | Location |
| --- | --- | --- | --- | --- | --- |
| CD14 | TUK4 | Qdot655 | Thermo Fisher | Q10056 | surface (pre-fixation) |
| CD45 | HI30 | 89Y | Fluidigm | 3089003B | surface |
| CD11c | Bu15 | 141Pr | Biolegend | 337221 | surface |
| CD11b | ICRF44 | 142Nd | Biolegend | 301337 | surface |
| CD45RA | HI100 | 143Nd | Biolegend | 304143 | surface |
| HLA-DR | L243 | 144Nd | Biolegend | 307651 | surface |
| CD4 | RPA-T4 | 145Nd | Biolegend | 300541 | surface |
| CD19 | HIB19 | 146Nd | Biolegend | 302247 | surface |
| CD20 | 2H7 | 147Sm | Biolegend | 302343 | surface |
| CCR6 | G034E3 | 148Nd | Biolegend | 353427 | surface (pre-fixation) |
| CD56 | NCAM16.2 | 149Sm | Fluidigm | 3149021B | surface (pre-fixation) |
| p-STAT5 | 47 | 150Nd | Fluidigm | 3150005A | intracellular |
| CD45RO | UCHL1 | 151Eu | Biolegend | 304239 | surface (pre-fixation) |
| CD27 | O323 | 152Sm | Biolegend | 302839 | surface (pre-fixation) |
| p-STAT1 | 4a | 153Eu | Fluidigm | 3153005A | intracellular |
| CD1c | L161 | 154Sm | Biolegend | 331502 | surface |
| CD123 | 6H6 | 155Gd | Biolegend | 306027 | surface |
| p-p38 | D3F9 | 156Gd | Fluidigm | 3156002A | intracellular |
| p-STAT3 | 4/P-Stat3 | 158Gd | Fluidigm | 3158005A | intracellular |
| p-MAPKAPK2 | 27B7 | 159Tb | Fluidigm | 3159010A | intracellular |
| CD3 | UCHT1 | 160Gd | Biolegend | 300443 | surface |
| DNGR1 | 8F9 | 161Dy | Fluidigm | 3161018B | surface |
| IFNAR2 | polyclonal | 162Dy | Abcam | ab56070 | surface |
| STAT1 | 246523 | 163Dy | Bio-Techne | MAB1490 | intracellular |
| IFNAR1 | EP899Y | 164Dy | Abcam | ab213331 | surface |
| CD161 | HP-3G10 | 165Ho | Biolegend | 339919 | surface |
| p-NFkBp65 | K10x | 166Er | Fluidigm | 3166006A | intracellular |
| CCR7 | G043H7 | 167Er | Fluidigm | 3167009A | surface (pre-fixation) |
| p-STAT6 | 18/P-Stat6 | 168Er | Fluidigm | 3168012A | intracellular |
| CD24 | ML5 | 169Tm | Fluidigm | 3169004B | surface |
| CD141 | M80 | 170Er | Biolegend | 344102 | surface |
| p-ERK1/2 | D13.14.4E | 171Yb | Fluidigm | 3171010A | intracellular |
| CD38 | HIT2 | 172Yb | Fluidigm | 3172007B | surface |
| STAT3 | 124H6 | 173Yb | Fluidigm | 3173003A | intracellular |
| p-STAT4 | 38/p-Stat4 | 174Yb | Fluidigm | 3174005A | intracellular |
| CCR4 | L291H4 | 175Lu | Fluidigm | 3175035A | surface (pre-fixation) |
| CXCR3 | G025H7 | 176Yb | Biolegend | 353733 | surface (pre-fixation) |
| CD8 | RPA-T8 | 198Pt | Biolegend | 301053 | surface |
| CD16 | 3G8 | 209Bi | Fluidigm | 3209002B | surface |

